# Inhibiting Glutamine Metabolism Blocks Coronavirus Replication in Mammalian Cells

**DOI:** 10.1101/2023.09.27.559756

**Authors:** Kai Su Greene, Annette Choi, Matthew Chen, Nianhui Yang, Ruizhi Li, Yijian Qiu, Michael J Lukey, Katherine S Rojas, Jonathan Shen, Kristin F Wilson, William P Katt, Gary R Whittaker, Richard A Cerione

## Abstract

Developing therapeutic strategies against COVID-19 has gained widespread interest given the likelihood that new viral variants will continue to emerge. Here we describe one potential therapeutic strategy which involves targeting members of the glutaminase family of mitochondrial metabolic enzymes (GLS and GLS2), which catalyze the first step in glutamine metabolism, the hydrolysis of glutamine to glutamate. We show three examples where GLS expression increases during coronavirus infection of host cells, and another in which GLS2 is upregulated. The viruses hijack the metabolic machinery responsible for glutamine metabolism to generate the building blocks for biosynthetic processes and satisfy the bioenergetic requirements demanded by the ‘glutamine addiction’ of virus-infected host cells. We demonstrate how genetic silencing of glutaminase enzymes reduces coronavirus infection and that newer members of two classes of small molecule allosteric inhibitors targeting these enzymes, designated as SU1, a pan-GLS/GLS2 inhibitor, and UP4, which is specific for GLS, block viral replication in mammalian epithelial cells. Overall, these findings highlight the importance of glutamine metabolism for coronavirus replication in human cells and show that glutaminase inhibitors can block coronavirus infection and thereby may represent a novel class of anti-viral drug candidates.

**Teaser:** Inhibitors targeting glutaminase enzymes block coronavirus replication and may represent a new class of anti-viral drugs.

## INTRODUCTION

Coronaviruses are a group of enveloped viruses that contain a single positive RNA strand and “corona”-like spike proteins extending from their envelopes. Seven types of coronaviruses have been reported to infect humans. In 2019, the Severe Acute Respiratory Syndrome 2 (SARS-CoV-2) virus triggered a worldwide pandemic, with multiple viral variants having emerged since that time. Human coronaviruses OC43 (HCoV-OV43) and 229E (HCoV-229E) are far less lethal and typically only give rise to common cold symptoms. Each coronavirus family member has a distinct spike protein and consequently binds to different receptors to enter cells. SARS-CoV-2 binds to the Angiotensin-converting enzyme 2 (ACE2) receptor^1–3^, HCoV-OC43 engages the N-acetyl-9-O-acetyl neuraminic acid receptor, and HCoV229E interacts preferentially with human aminopeptidase (hAPN)^2–7^. SARS-CoV-2 and HCoV-OC43 are members of the coronavirus beta sub-group family, while HCoV-229E is a member of the alpha sub-group. Since the onset of the pandemic, therapeutic antibodies and vaccines targeting SARS-CoV-2 have been developed, although each has its limitations. Thus, there continues to be a pressing need to identify new therapeutic strategies that will provide broad protection against new virus strains and mutants which are likely to emerge.

Viruses have been suggested to reprogram the metabolism of host cells to support their bioenergetic requirements for replication, raising the possibility that targeting metabolic activities essential for viral infections might offer potential therapeutic strategies. For example, there have been reports suggesting that some viruses are dependent upon glutamine metabolism for their replication and the generation of new viral particles including Kaposi-sarcoma associated herpesvirus (KSHV), vaccinia virus (VACV), adenovirus (AD) and influenza A virus (IAV)^8–9^. Normal healthy cells typically utilize glucose as their primary bioenergetic and biosynthetic material, whereas, cancer cells often reprogram their metabolism by increasing the utilization of glutamine as the primary nutrient to support their TCA cycle and satisfy the metabolic requirements necessary for their high rates of proliferation and ability to survive various types of cellular stress^10–15^. Members of the glutaminase family of enzymes play a critical role in satisfying these metabolic requirements by catalyzing the first step in glutamine metabolism, the hydrolysis of glutamine to glutamate, with glutamate then being converted to α-ketoglutarate by glutamate dehydrogenase to enter the TCA cycle.

Two genes, *Gls* and *Gls2*, encode the glutaminase enzymes in mammals^15^. *Gls* encodes KGA (kidney-type glutaminase) and the C-terminal truncated splice variant GAC (glutaminase C), hereon collectively referred to as GLS, which are ubiquitously expressed in mammalian tissues, while *Gls2* encodes LGA (liver-type glutaminase, hereon designated GLS2) and is primarily expressed in liver, pancreas, and brain^16–19^. GLS has been shown to be highly expressed in various types of cancers including basal-subtype triple negative breast cancer, glioblastoma, and pancreatic cancers^20,21^, due to the actions of the transcription factors c-Myc and c-Jun^22,23^ GLS2 was recently implicated in luminal-subtype breast cancer^21,24^. We and others have reported that allosteric inhibitors of GLS, including 968, BPTES, CB839, and UPGL00004 (designated as UP4 from hereon) block cancer cell proliferation^18^_’_^25^, with CB839 being examined in a number of clinical trials as an anti-cancer drug^19,26,27^.

GLS inhibitors including BPTES and CB839 have also been shown to block the replication of some viruses^8,9^. However, thus far, it has not been demonstrated that glutamine metabolism is essential for coronavirus infection and if GLS inhibitors might be effective in blocking their replication^28^. Here we show that three members of the coronavirus family, SARS-CoV-2, HCoV-OC43, and HCoV-229E reprogram the metabolic machinery of host cells, and in doing so, cause their replication to become glutamine-addicted. In these studies, we have used different cell lines for each coronavirus based on the expression of viral binding receptors; the mouse kidney epithelial cell line VeroE6 for SARS-CoV-2 infections, the human bronchial epithelial cell line HBEC and human colon cancer HCT8 epithelial cells for HCoV-OC43, and the lung epithelial cell line MRC5 for HCoV-229E (Table 1). We then demonstrate that glutaminase inhibitors, including two new compounds (SU1 and UP4) that we have developed, represent potential therapeutic agents for blocking coronavirus infection with broad application.

**Table 1.**
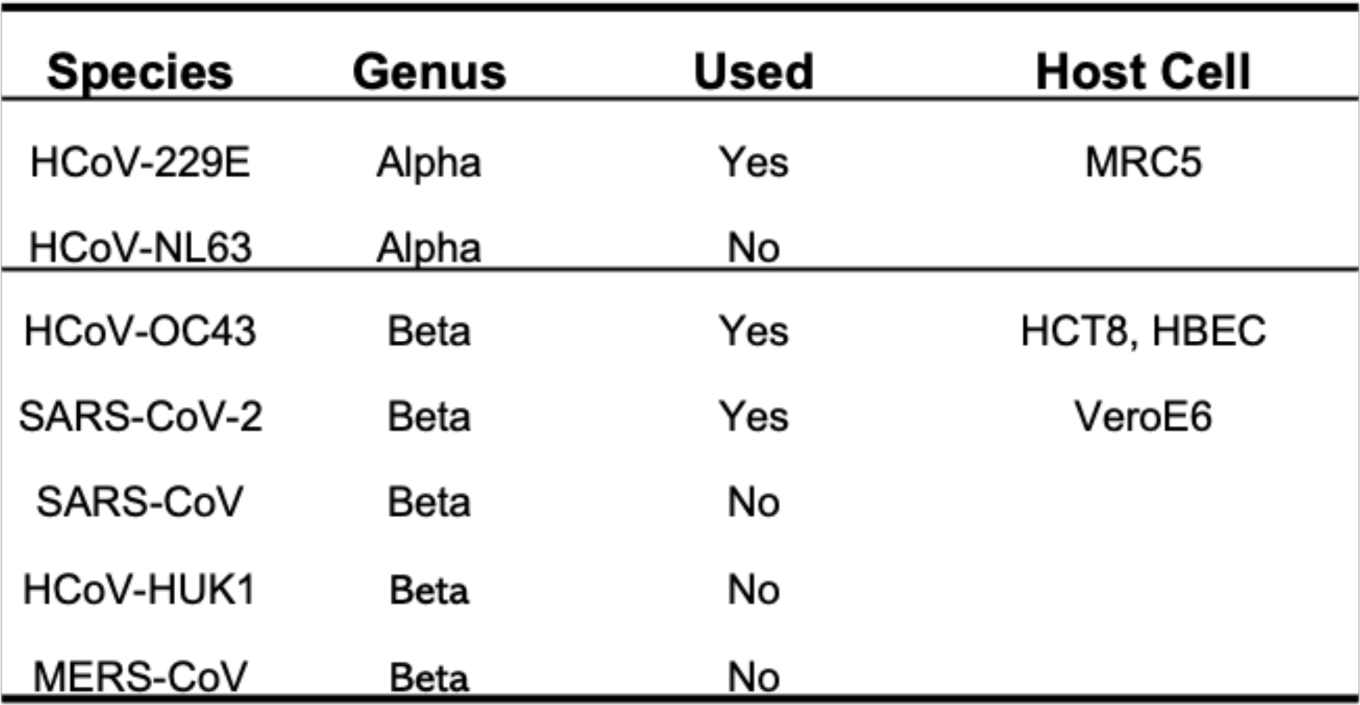
Human coronaviruses and host cells. There are seven human coronaviruses. The three viruses used for this study are listed with their host cells.

## RESULTS

### Coronavirus infection induces metabolic reprogramming in host cells

Viruses induce metabolic reprogramming in infected cells, and their replication depends on biosynthetic precursors and ATP provided by the host^29^. Since glutamine supplies multiple biosynthetic pathways with carbon and/or nitrogen^8,30^, we were interested to see whether coronavirus infection upregulates glutamine metabolism in primary human bronchial epithelial cells HBEC3-KT (HBEC from hereon). We therefore applied LC-MS-based targeted metabolomics to metabolite extracts from HBECs that were either uninfected or infected for 24 hours with the human beta-coronavirus HCoV-OC43. Upon infection, there was an apparent activation of glutamine metabolism, as read-out by increases in the products of glutaminolysis, namely glutamate and α-ketoglutarate, and downstream metabolites such as fumarate and aspartate (Figs. 1A and 1C). We also observed an increase in the levels of several nucleotides in infected cells, consistent with the known dependence of viral replication on activated nucleotide biosynthesis in host cells^31^ (Fig. 1B).

**Figure 1.**
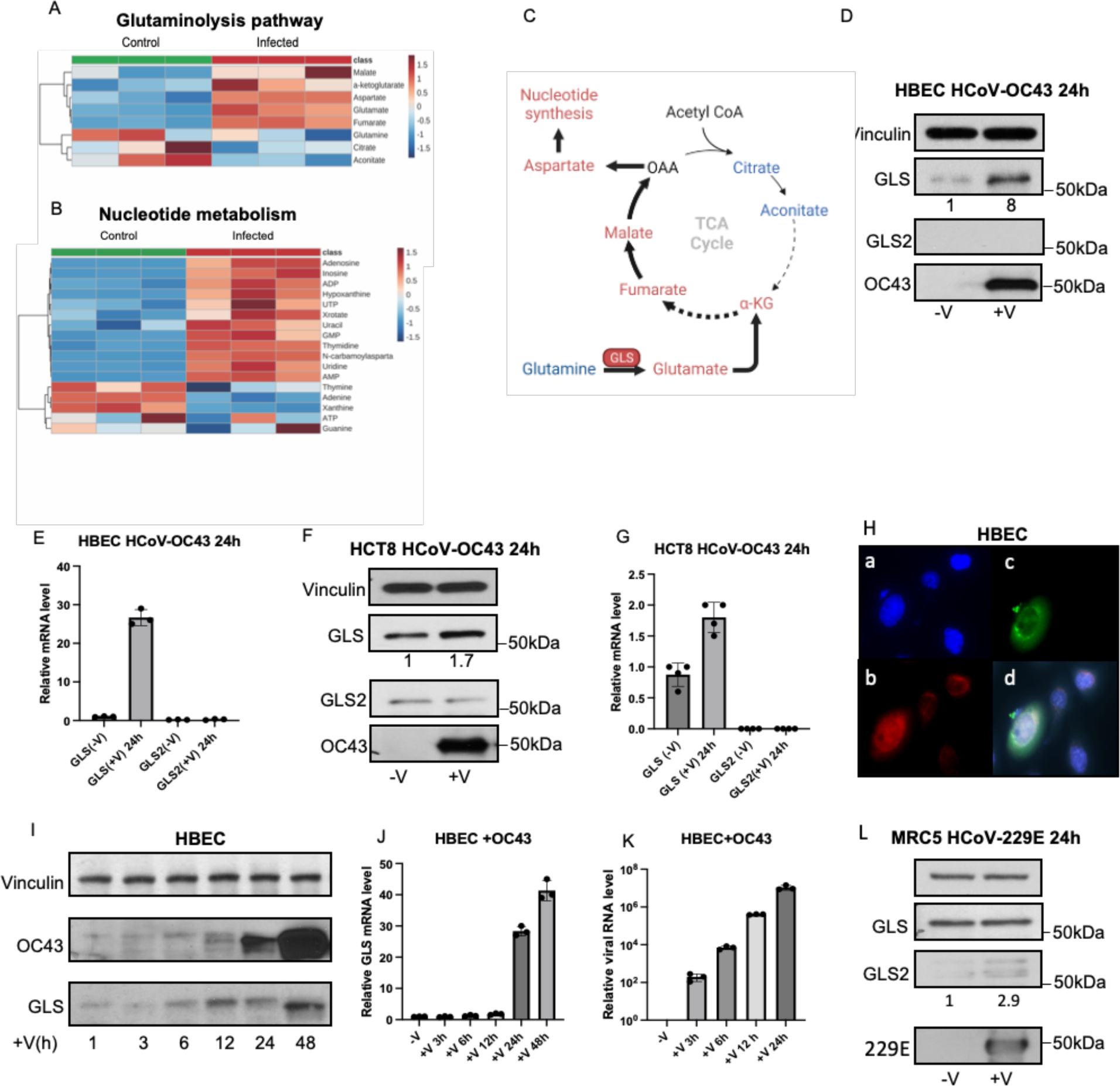
GLS expression is increased during coronavirus infection. (A) HBECs were infected with HCoV-OC43 (MOI 0.01) for 24 hours before metabolites were extracted. Related carbon metabolic pathways, including TCA cycle, glycolysis, pentose phosphate pathway, and nucleotide synthetic pathways, were analyzed via targeted metabolomics. The heatmap showed the Glutaminolysis pathway components change before and after virus infection. (B) Heatmap of Nucleotide metabolism components change before and after virus infection. (C) The diagram shows the upregulated (in red) and downregulated (in blue) metabolites in HCoV-OC43-infected HBECs. (D) Whole cell lysates of uninfected HBECs or HBECs infected with HCoV-OC43 (MOI 0.01, 24 h) were analyzed for viral N-protein OC43, GLS and GLS2 expression levels by Western blot. (E) Total RNA isolated from uninfected HBECs or HBECs infected with HCoV-OC43 (MOI 0.01, 24 h). qPCR assays showed GLS and GLS2 mRNA levels before and after HCoV-OC43 infections. (F) Similar experiment as (D) in HCT8 cells. (G) qPCR assays performed as (E) in HCT8 cells. (H) Co-immunofluorescent staining with HCoV-OC43 and GLS antibodies in HBECs. Cells were grown in 4-well slide chamber for 24 hours and were infected with the HCoV-OC43 virus for 24 hours, One drop per well of NucBlue Live Cell stain solution was added to each well before the cells were fixed with formaldehyde (3.7%) (a) nuclear staining with blue color, (b) staining with GLS rabbit polyclonal antibody and Goat anti-rabbit Alexa Fluor^TM^ 568 Secondary antibody, (c) staining with HCoV-OC43 mouse monoclonal antibody and Goat anti-mouse Alexa Fluor^TM^ 488 Secondary antibody. (d) Merge of the images described in a,b,c. (I) Western blot analysis shows the time course of GLS and HCoV-OC43 expression levels during virus infection in HBECs. (J) qPCR assays were performed on total RNA samples isolated at different time points of virus infected HBECs. (K) qPCR assays were performed on the total RNA samples isolated at different time points of virus-infected media in HBECs. (L) Whole cell lysates of MRC5 cells uninfected or infected with HCoV-229E (MOI 0.01, 24h) were analyzed for GLS and GLS2 expression levels by Western blot. (M) qPCR assays for GLS2 transcript levels were performed with the total RNA isolated from uninfected or HCoV-229E infected MRC5 cells.

### Glutaminase expression is upregulated during coronavirus replication

Based upon the changes in glutamine metabolism caused by coronavirus infection, we examined whether glutaminase expression was upregulated when host cells were infected with HCoV-OC43. HBECs and HCT8 cells, when 80% confluent, were infected with HCoV-OC43 for 24 hours, and then the cells and their medium were collected. Western blot analyses were performed using an anti-HCoV-OC43 antibody to detect the viral nucleoprotein OC43 as a read-out for viral replication, and anti-GLS and anti-GLS2 antibodies were used to identify the two forms of glutaminase expressed in the human epithelial cells (Figs. 1D and 1F). Although GLS protein expression was relatively low in HBECs, it increased significantly (eightfold) upon HCoV-OC43 infection. On the other hand, GLS2 protein expression was not detected before or after virus infection, indicating that only GLS is involved in coronavirus HCoV-OC43 replication in HBECs. In HCT8 cells, GLS protein expression was modestly increased (1.7 fold) with virus infection, whereas GLS2 protein levels were not changed (Fig. 1F). However, the basal levels of both GLS and GLS2 were higher in HCT8 cells compared to HBECs, most likely because the former represents a human colon cancer cell line. Quantitative PCR (qPCR) assays showed that the RNA transcript levels of GLS but not GLS2 were upregulated in both virus-infected HBECs and HCT8 cells (Figs. 1E and 1G).

We then carried out immunofluorescence experiments in HBECs to visualize GLS expression as a function of virus infection at a cellular level. Cells were cultured in a 4-well chamber slide with 60-70% confluency, infected the next day with HCoV-OC43 as described above for 1 hour, and then incubated in fresh culture medium for 22 hours with and nuclear staining for 1 hour with NucBlue. After fixation, immunofluorescence staining was carried out using antibodies specific for either GLS or the HCoV-OC43 N protein, with the cell nuclei being visualized. Four HBECs shown in Figure 1H-a (blue) exhibited different levels of GLS (Fig. 1H-b, red) and HCoV-OC43 expression (Fig. 1H-c, green). The merged images show that the virus-infected cells consistently expressed higher levels of GLS (Fig. 1H-d).

Next, we performed a time course for HCoV-OC43 infection of HBECs. The cells (70-80% confluent) were infected with HCoV-OC43 for 1 hour, with the cells from one plate then being collected. Fresh medium was added to the remaining plates, and additional samples of cells and their culture medium were collected after 3, 6, 12, 24, and 48 hours of incubation at 37°C. The expression levels of the HCoV-OC43 nucleoprotein OC43 and GLS were detected by Western blot analyses from whole cell lysates (Fig. 1I), while qPCR was used to measure the total RNA transcript levels of GLS (Fig. 1J) and of viral RNA in the media (Fig. 1K). OC43 and GLS expression levels in HBECs showed significant increases between 12 and 48 hours of virus infection. The increase in GLS expression followed the time course for that of the virus protein OC43 (Fig. 1I), and the same was true for their RNA transcript levels (Fig. 1J), indicating that the upregulation of GLS expression began with the onset of virus infection.

We also examined glutaminase expression in MRC5 cells infected by HCoV-229E (an alpha coronavirus). MRC5 cells at 80% confluency were infected with HCoV-229E (MOI 0.01, 1 hour, at 33°C), and then incubated for 23 hours with growth medium at 37°C. Western blot analyses of whole cell lysates showed that GLS2 expression was increased but not GLS (Fig. 1L).

### GLS is essential for coronavirus replication in HBECs

To determine whether GLS expression is necessary for coronavirus infection, we knocked down GLS in HBECs for 24 hours, using two different shRNAs. The cells were then infected with HCoV-OC43 for 24 hours, and cell lysates were analyzed by Western blotting to determine GLS expression, while their medium was collected and analyzed by qPCR to detect total viral RNA. We found that HCoV-OC43 replication was reduced significantly when GLS was depleted from the cells, as evidenced by the decreased expression of the viral protein OC43 (Fig. 2A) and the reduced amount of viral RNA (72% and 80% respectively with two shRNAs) and in the level of viral RNA transcripts in the medium (Fig. 2B). Similar reductions in HCoV-OC43 protein expression (Fig. 2C), and in the levels of viral RNA transcripts in the medium (Fig. 2D) were observed when using two independent siRNAs targeting GLS.

**Figure 2.**
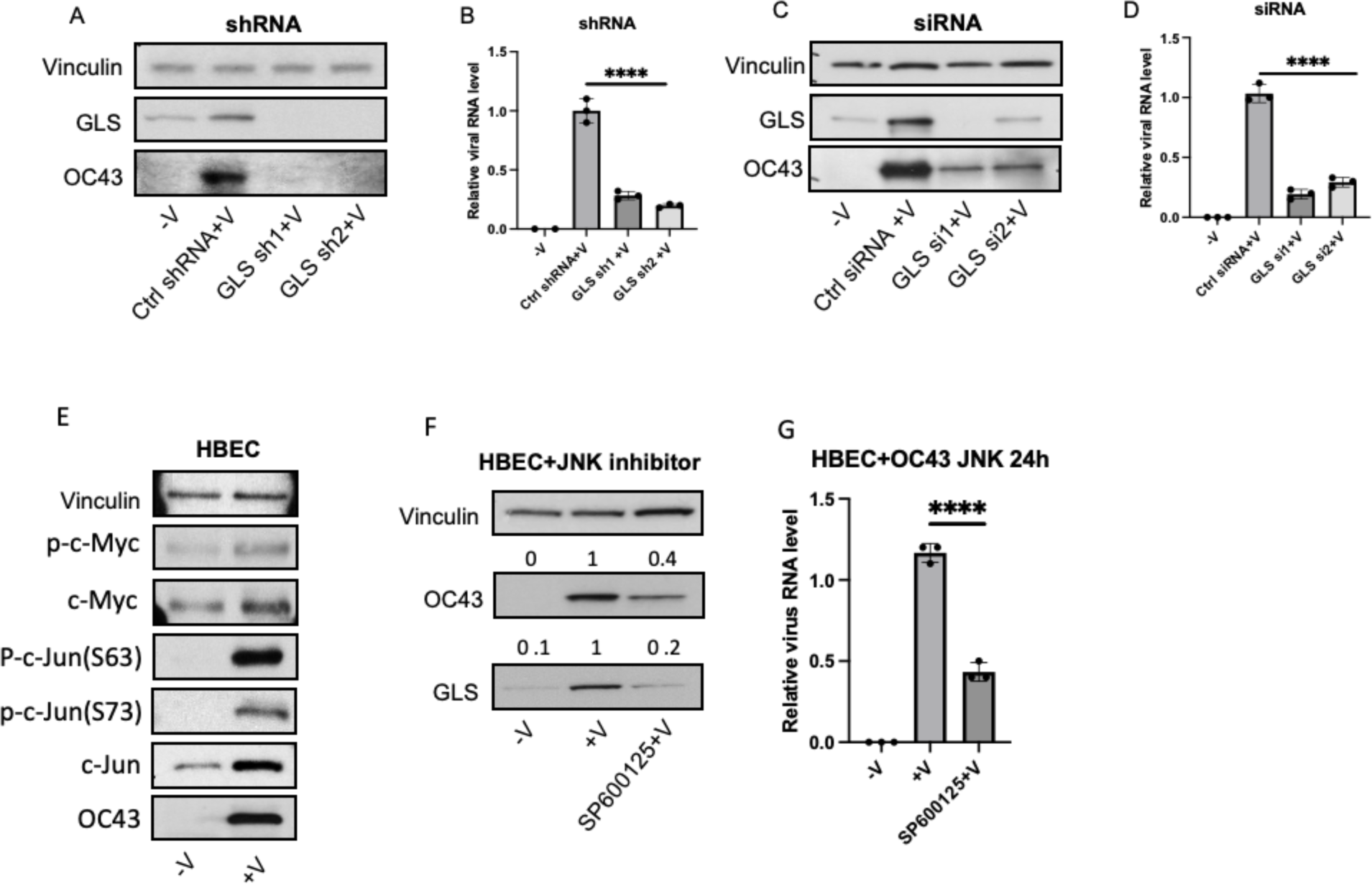
Glutaminase is an essential enzyme for coronavirus replication. (A) Western blot analysis showing HCoV-OC43 replication levels in HBECs expressing a control shRNA or two independent GLS-targeted shRNAs. (B) qPCR assays showing the relative virus RNA levels in the media of control and two GLS-targeted shRNAs. (C) Western blot analysis shows HCoV-OC43 replication levels in HBECs expressing control or two GLS-targeted siRNAs. (D) qPCR analysis showing the total virus RNA levels from the media of non-virus infection, virus-infected cells with control siRNA, and two GLS-targeted siRNA knockdowns. (E) HBECs were infected with HCoV-OC43 for 24 hours (described previously) with the non-treated cells as a control. The whole cell lysates were collected. Western blot analysis shows that p-c-Jun(S73), p-c-Jun(S63), c-Jun, p-c-Myc and c-Myc levels are upregulated upon HCoV-OC43 infection of HBECs. (F) Western blot data showing that blocking c-Jun activation by inhibiting the c-Jun-N-Terminal kinase (JNK) with inhibitor SP600125 (10µM) for 24 hours reduces GLS expression in virus-infected host cells. (G) qPCR assays showing the relative HCoV-OC43 mRNA levels in the media from cells treated with or without JNK inhibitor. One-way ANOVA with Bonferroni correction was used to determine significance in C, **** indicates modified *P* < 0.0001.

### GLS expression is upregulated in HBECs by c-Jun during virus infection

GLS expression in cancer cells has been reported to be upregulated either by c-Myc or c-Jun^22,23^, and the JNK/c-Jun pathway has been shown to be activated in some infected cells^32–35^. To further examine how GLS expression is increased upon virus infection, HBECs were infected with HCoV-OC43 for one hour followed by a 23-hour incubation, at which point the cells were collected and Western blot analyses performed. Phospho-c-Myc, phospho-c-Jun(S63) and phospho-c-Jun(S73) were all observed to be increased after infection, as were the total protein expression levels for c-Myc and c-Jun together with the viral protein OC43 (Fig. 2E). However, while inhibiting c-Myc did not cause a reduction in the expression levels of GLS in virus-infected host cells (HCT8) (not shown), blocking c-Jun activation using the small molecule inhibitor SP600125 reduced both GLS protein expression and the amount of viral RNA transcripts detected in the medium (Figs. 2F and 2G, respectively)

### Glutaminase inhibitors block coronavirus SARS-CoV-2, HCov-OC43 and HCoV-229E replication

We next examined whether small molecule allosteric inhibitors targeting the glutaminase enzymes could impact coronavirus infection. We and others have developed and characterized two classes of allosteric glutaminase inhibitors, based on the lead compounds 968 and BPTES^20,36–39^ (Fig. 3A). X-ray crystal structures have shown that BPTES, and its more potent analogs CB839 and our newly developed inhibitor UP4, bind in the interface where two dimers of GLS come together to form a tetramer ^40^, while 968 and the more potent SU1 compound^33^ have been suggested to bind in close proximity but not directly overlapping the BPTES/CB839/UP4 binding sites (Fig. 3B)^45^. Both classes of inhibitors trap the enzyme in an inactive tetrameric state, although the 968 group of compounds also induce the formation of some inactive dimer species^45^. In these experiments, we examined HCoV-229E which belongs to the alpha-genus of the coronavirus family, and SARS-CoV-2 and HCoV-OC43 from the beta-genus (Table 1).

**Figure 3.**
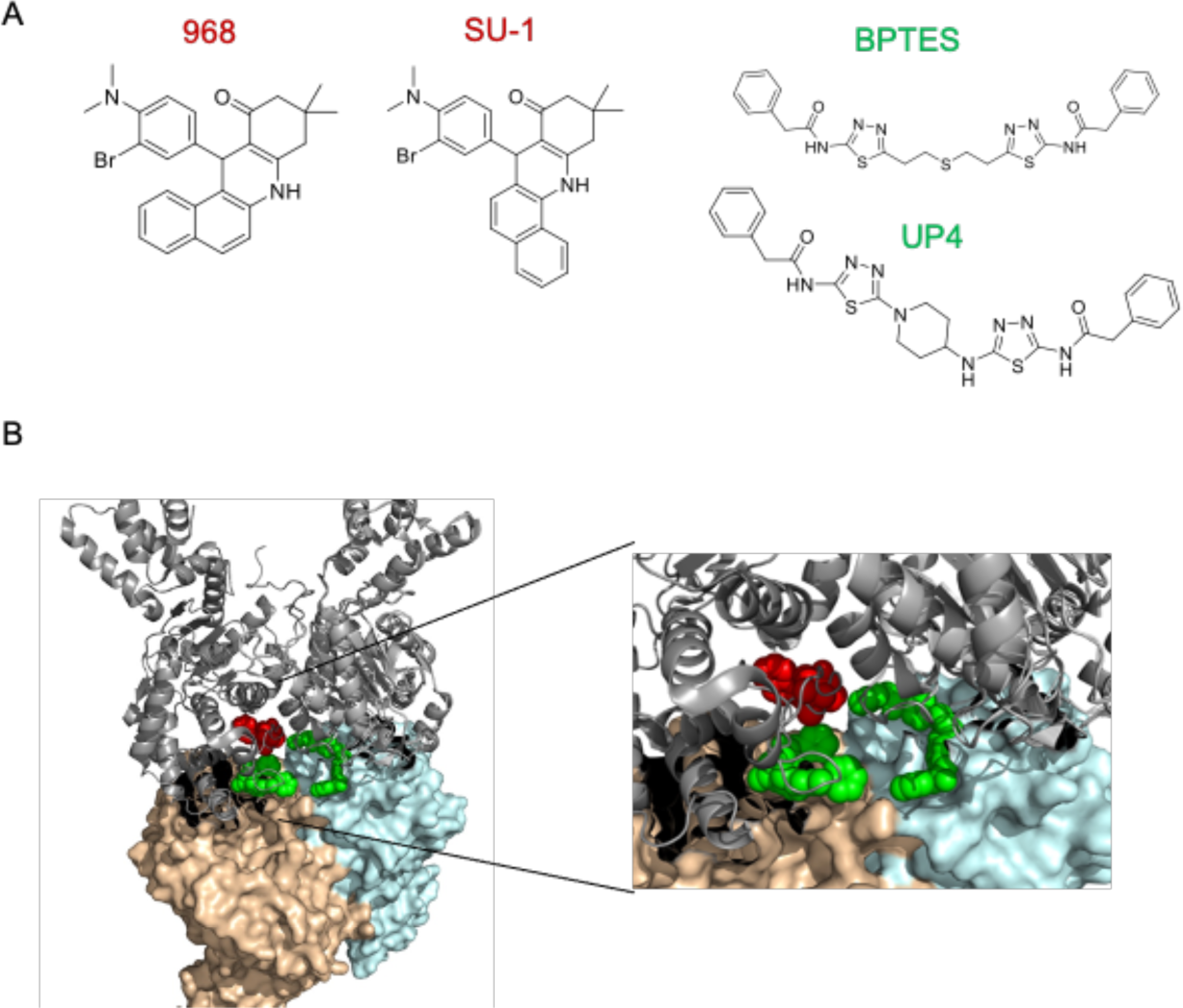
Allosteric inhibitors targeting GLS. (A) The structures of two classes of glutaminase inhibitors: 968 and the more potent analog SU1; and BPTES and its more potent analog UP4. (B) Homo-tetramer of human GLS structure showed the proposed binding site for 968 and SU1 (red color) based on a recent cryo-EM structure for the 968-GLS complex and the binding location for BPTES and UP4 class (green) as determined by X-ray crystallography.

The experiments were started by growing cells to 70-80% confluence and pretreating for 3 hours with glutaminase inhibitor or vehicle control, followed by infection of VeroE6 cells with SARS-CoV-2 (MOI 0.01, 2%FBS in DMEM) for 1 hour at 37°C, and HBECs or HCT8 cells with HCoV-OC43, (MOI 0.01, 2% FBS in RPMI medium) for 1 hour at 33°C. We tested two groups of glutaminase inhibitors, the 968 analog and pan-GLS/GLS2 inhibitor SU1, and the BPTES related compound and GLS selective inhibitor UP4. After a 23-hour incubation at 37°C, the cells and their media were collected, and Western blot analyses were performed on the cell lysates to detect the total levels of the SARS-CoV-2 (Fig. 4A) and HCoV-OC43 proteins (Figs. 4B and 4C). We observed that virus replication of SARS-CoV-2 and HCoV-OC43 was significantly suppressed by the glutaminase inhibitors in a dose-dependent manner, as read-out by reductions in the expression of the SARS-CoV-2 spike protein and the HCoV-OC43 protein (Figs. 4A-C, and Figs. S3A-D). We confirmed these results by plaque assays (Figs. S4A and S4B) and by qPCR measurements of the virus RNA levels for SARS-CoV-2 in the VeroE6 medium (Fig. 4D) and the HCoV-OC43 RNA levels in both HBEC (Fig. 4E) and HCT8 media (Fig. 4F).

**Fig. 4.**
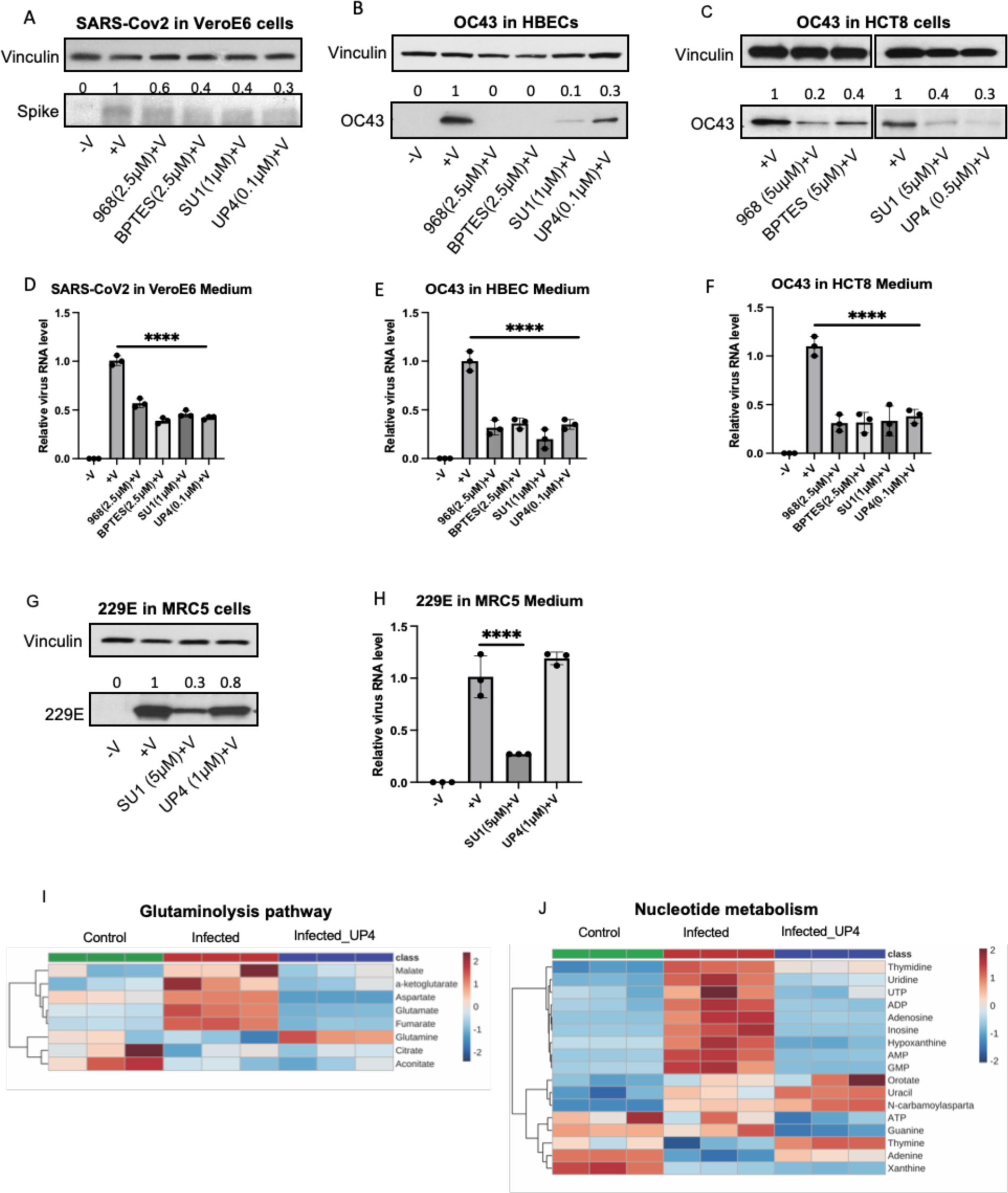
GLS inhibitors block coronavirus SARS-CoV-2, HCoV-OC43 and HCoV-229E replication. (A) VeroE6 cells when 70-80% confluence were pretreated with 968 (2.5μM), BPTES (2.5μM), SU1 (1μM), UP4 (0.1μM), and vehicle control (DMSO) for 3 hours, followed by SARS-CoV-2 infection (MOI 0.01) for 1 hour at 37°C. The growth medium was changed with or without inhibitors and the cells were incubated at 37°C for 23 hours. The media and cells were collected. Western blot analysis of the inactivated whole cell lysates showing the SARS-CoV-2 replication levels in cells treated with the different inhibitors compared with DMSO treated virus-infected cells. (B) HBECs were pretreated with DMSO, 968 (2.5μM), BPTES (2.5μM), SU1 (1μM), and UP4 (0.1μM) for 3 hours, followed by HCoV-OC43 infection for 1 hour at 33°C, and then cultured in growth medium with or without inhibitors for 23 hours at 37°C. Western blot analysis shows the OC43 replication levels in cells treated with different inhibitors compared with no drug treatment. (C) HCT8 cells were infected with HCoV-OC43 as in (B) and analyzed for virus replication by Western blotting for OC43. (D) qPCR assays showing the relative virus RNA levels in the SARS-CoV-2 infected cell media. (E) qPCR analysis showing the OC43 virus RNA levels in the media of HCoV-OC43 infected HBECs with or without inhibitors. (F) HCoV-OC43 virus RNA levels from the media of HCT8 cells treated with or without inhibitors. (G) MRC5 cells were pretreated with DMSO, SU1 (2.5µM) and UP4 (0.5µM) for 3 hours, followed by HCoV-229E infection for 1 hour at 33° C. The growth medium was changed to that for the pretreated condition for 23 hours and then the cells and medium were collected. Western blot assays of the cell lysates showed the 229E virus levels in different conditions. (H) qPCR assay showing the relative viral-RNA levels of MRC5 infected with 229E media with or without GLS inhibitors. (I) HBECs pre- and post-treated with UP4 (0.5µM) were infected with HCoV-OC43 (MOI 0.01) for 24 hours before metabolites were extracted. Related carbon metabolic pathways were analyzed via targeted metabolomics. The heatmap showed Glutaminolysis pathway component changes with or without UP4 treatment during virus infection compared with wildtype cells. (J)The heatmap Nucleotide metabolism components before and after virus infection or virus infection treated with UP4. One-way ANOVA with Bonferroni correction was used to determine significance in D, E, F, **** indicates modified *P* < 0.0001.

Similar experiments were performed with MRC5 cells infected with HCoV-229E. MRC5 cells were pretreated with DMSO, SU1 (2.5µM), UP4 (0.5 µM) for 3 hours, and then the cells were infected with HCoV-229E (MOI 0.01, 1 hour at 33°C), followed by an incubation at 37°C with fresh culture medium for 23 hours. The cells and media were collected and then Western blot and qPCR analyses performed. The data indicate that in MRC5 cells, the pan-GLS/GLS2 inhibitor SU1 blocks HCoV-229E replication by inhibiting GLS2 activity, whereas the GLS-specific inhibitor UP4 was ineffective (Figs. 4G and 4H).

### Glutaminase inhibition reverses coronavirus induced metabolic reprogramming

We next examined how inhibiting glutaminase activity affects coronavirus-induced metabolic reprogramming in host cells. HBECs were infected with HCoV-OC43, whose viral replication was sensitive to both classes of glutaminase inhibitors. The cells were infected for 24 hours in the absence or presence of 0.5 µM UP4, and cell extracts were then prepared for targeted metabolomics analysis. Heatmaps of Glutaminolysis pathway and Nucleotide metabolism of the three groups of HBECs, non-infected, infected with HCoV-OC43, or infected with HCoV-OC43 in the presence of UP4 (Figs. 4I and 4J). As expected, treatment with UP4 abolished the increases in glutamate, downstream TCA cycle intermediates, and aspartate during OC43 infection (Fig. 4I), as well as those of nucleosides and nucleotides (Fig. 4J) and similar results were obtained with SU1 (Fig. S5).

### Glutaminase inhibitors do not block coronavirus entry into host cells

The entry of coronaviruses into their host cells are dependent upon binding to membrane-associated receptors. HBECs lack the ACE2 receptor and thus we did not observe these cells to be infected by SARS-CoV-2. However, HCoV-OC43 can infect HBECs as well as HCT8 cells (Figs. 1D, 1E, 1F, 1G and 1L). We then tested whether GLS inhibitors block coronaviruses from entering cells. HBECs and HCT8 cells were pretreated with either SU1 or UP4 or untested, for 3 hours. The cells were then infected with HCoV-OC43 (MOI 0.01, 2%FBS in RPMI) for 1 hour at 33°C, at which point they were collected, and Western blot analyses were performed to detect the expression of the OC43 viral protein (Fig. 5A). We found that OC43 was present under all conditions of inhibitor treatments. The presence of viral RNA in the cells as determined by qPCR was also unaffected by the inhibitors (Fig. 5B). Thus, taken together, these findings indicate that glutaminase inhibitors block coronavirus infection by preventing viral replication rather than by inhibiting viral entry.

**Figure 5.**
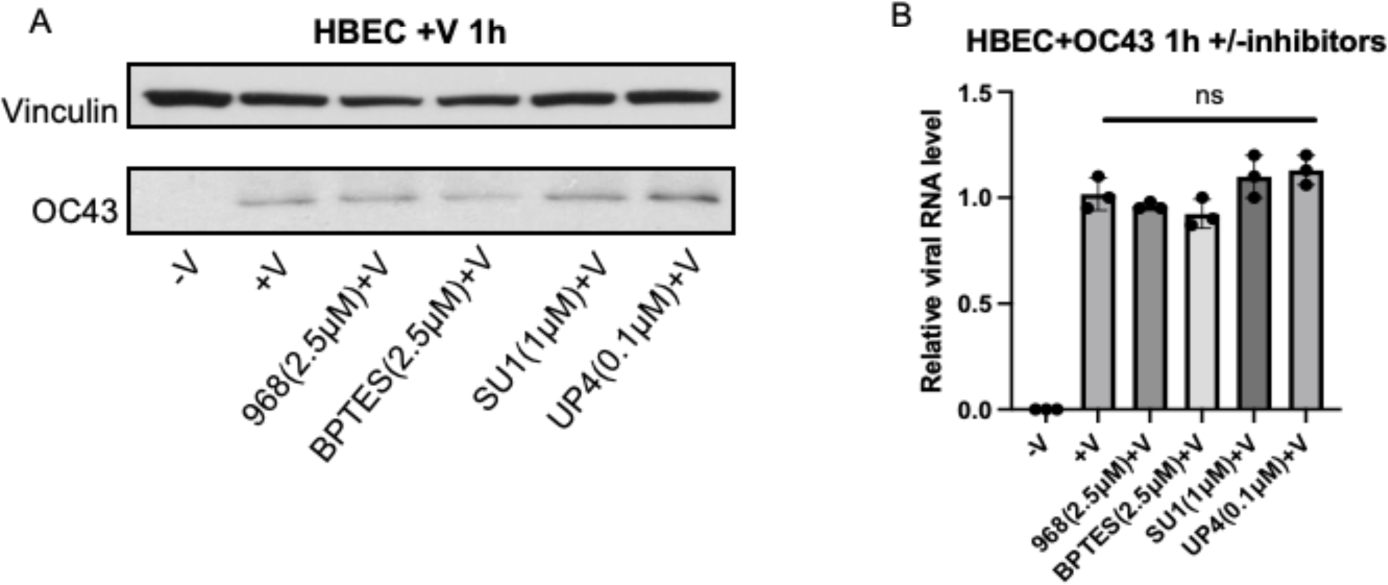
GLS inhibitors do not block coronavirus entry. (A) HBECs were pre-treated with DMSO, 968(2.5μM), BPTES(2.5μM), SU1(1μM), and UP4(0.1μM) for 3 hours, and then the cells were infected with HCoV-OC43 (MOI 0.01) for 1 hour at 33°C 5%CO_2_ and collected. Western blot analysis showing the HCoV-OC43 levels in the HBECs with or without inhibitors. (B). HBECs were treated with inhibitors and infected with the virus as above. The total viral RNA levels for each condition were analyzed by qPCR.

## DISCUSSION

The glutaminase family of mitochondrial metabolic enzymes has been shown to play important roles in cancer progression due to the ability of its members to catalyze the first step in glutamine metabolism, the hydrolysis of glutamine to glutamate with the accompanying production of ammonia^11,13,16,18,23,41^. By increasing glutaminolysis, cancer cells satisfy their metabolic requirements and glutamine addiction, which are an outcome of the Warburg effect in which the glycolytic pathway is uncoupled from the TCA cycle. The elevations in glutamine metabolism that occur in cancer cells provide the carbon sources necessary to generate building blocks for biosynthetic processes that underlie their malignant phenotypes. It has been reported that some virus-infected host cells appear to undergo a reprogramming of their metabolism similar to cancer cells^21,35^. Here we show that this is the case for coronavirus infection including SARS-CoV-2 by demonstrating that glutamine metabolism in the host cells is essential for their replication.

There are two major forms of the glutaminase enzymes in mammals and humans, designated here as GLS and GLS2. Both GLS expression and its specific activity has been implicated in a number of human cancers, although GLS2 has also been shown to be important in luminal-subtype breast cancer^21^. In our studies, we have examined four different host cell lines and three members of the coronavirus family and found that in most cases GLS is essential for viral replication; however, in one host cell, MRC5, it appears that GLS2 is the glutaminase enzyme required for viral replication. The upregulated expression of GLS in cancer cells has been shown to be an outcome of either c-Myc blocking the inhibitory actions of a microRNA^22^, or through signaling pathways that result in the activation of the transcription factor c-Jun^23^. For host cells infected by coronaviruses, we have found that it is c-Jun activation that upregulates GLS expression. Thus far, very little is known regarding how glutaminase activity is activated either in cancer or virus-infected cells. We and others have shown that both GLS and GLS2 activation requires that the enzyme undergoes a transition from an inactive dimer to a tetramer and then ultimately to a higher-order filament^44,45^. The formation of activated glutaminase filaments requires both the binding of substrate and an anionic activator. Inorganic phosphate is commonly used to serve as an activator *in vitro*, although the concentrations required (50-100 mM) are unlikely to be achieved in most physiological settings and therefore, we are setting out to determine what type of metabolite or cofactor might serve this function in cells.

There has been a significant amount of effort devoted to developing small molecule inhibitors that target the glutaminase enzymes, given their roles in tumorigenesis. Two of the more common types of GLS inhibitors are the 968 class of molecules and the BPTES family of inhibitory compounds^11,14,17,39,47^. Among the latter are CB839 and our more newly developed UP4^40^, which are significantly more potent than the lead compound BPTES, with CB839 being examined in clinical trials for various cancers^41,48,49^. The BPTES series of compounds bind within the interface where two GLS dimers come together to form a tetramer and stabilize an inactive tetrameric species that is incapable of forming higher order filament-like structures^45^. The 968 class of GLS inhibitors which includes the lead compound 968 and our more potent analog SU1^38^ function in a distinct manner from the BPTES class of compounds, by preferentially binding initially to GLS monomers and then stabilizing both inactive dimers and tetramers. Both classes of allosteric GLS inhibitors effectively blocked coronavirus replication in HBECs, HCT8 and VeroE6 cells, as read out by the synthesis of a viral coat protein and when assaying viral RNA transcript levels, matching the effects we observed when knocking down GLS expression in host cells. However, in MRC5 cells, only the 968 analog SU1 was effective at inhibiting viral replication because of the required role of GLS2, which is relatively insensitive to UP4 and other related compounds^21^. These findings are summarized in the schematic presented in Figure 6.

**Figure 6.**
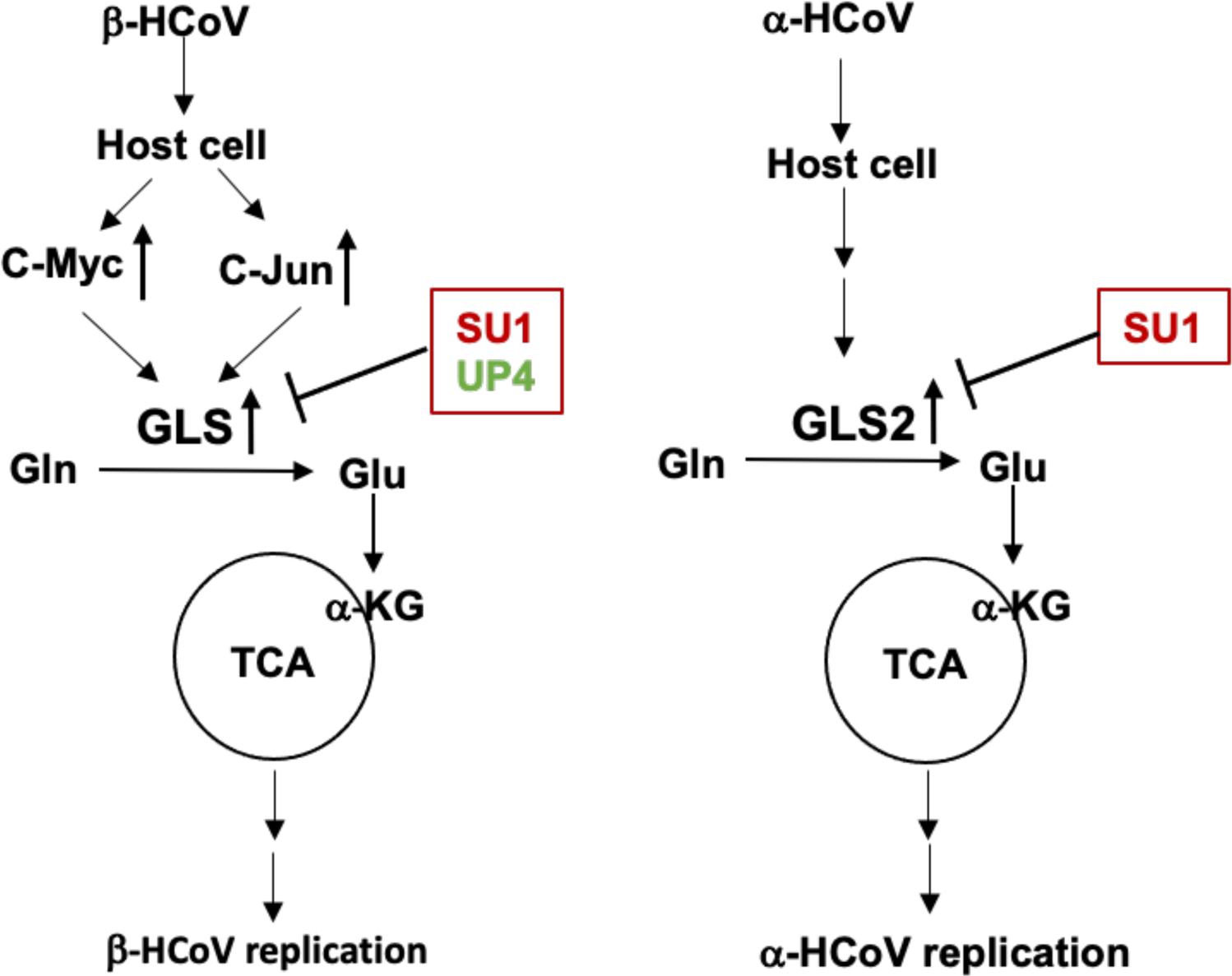
Schematic of GLS mediated coronavirus replication. Coronaviruses hijack the metabolic program of host cells by increasing the expression of GLS. Two possible mechanisms for upregulating GLS upon beta-coronavirus infection involve signaling through c-Myc and c-Jun. Alfa-coronavirus replication occurs through the upregulation of GLS2 expression (HCoV-229E in MRC5 cells).

Current approved anti-coronavirus drugs target proteins essential for virus replication, including RNA polymerases^50–52^, spike proteins^53,54^, proteases such as TMPRSS2^4,54^, and nucleocapsid proteins^55^. Our work has focused on blocking the metabolic activities required to generate building blocks for biosynthetic processes and the energy supply necessary for coronavirus replication. Because the glutaminase enzymes are often much more highly expressed and activated in cancer^5,6,8,45^ and viral-infected host cells^8,9,30,57^, as compared to normal healthy cells, small molecule inhibitors targeting these enzymes offer a potentially safe therapeutic strategy. Here we now show that three coronaviruses including SARS-CoV-2 require glutamine metabolism for their replication and are susceptible to inhibition by allosteric glutaminase inhibitors. We demonstrate how glutaminase inhibitors are especially effective at blocking coronavirus infection and that they may serve as new lead compounds toward the design of even more potent and novel anti-viral drug candidates.

Finally, our findings raise interesting questions and lines of investigation for future studies. They include determining how virus infection triggers the necessary signals to upregulate glutaminase expression and elucidating the mechanism responsible for the activation of GLS catalytic activity. It also will be of interest to establish whether glutaminase filament-like structures form within virus-infected host cells, similar to the case for cancer cells^44,45,58,59^. If so, do these higher order oligomeric structures serve as a scaffold for a metabolic complex necessary for satisfying the requirements for viral replication? Recently we showed that glutaminase inhibitors can block the ability of both GLS and GLS2 to form filaments^45^ and it will be interesting to see if additional strategies be designed to specifically block their formation and thus yield additional classes of anti-viral therapeutics.

## MATERIALS and METHODS

### Materials

#### Coronaviruses

SARS-CoV-2 WA/1 (BEI resources, cat. NR-52281), HCoV-OC43 (Betacoronavirus 1, cat. VR-1558), and HCoV-229E (Human coronavirus 229E, cat. VR-740) were from ATCC.

#### Cell Lines

HCT8 (human colorectal carcinoma cell line initiated from an adult male, cat. CCL-244), HBEC3-KT (Primary human bronchial epithelial cells, cat. CRL-4051), MRC5 (normal human fetal lung fibroblast cells, cat. CCL-171), and VeroE6 (Kidney epithelial cells isolated from African green monkey, cat. CRL-1587) were all from ATCC.

#### Antibodies

The GLS antibody was raised against the sequence-KLDPRREGGDQRHS; GLS2 antibody was obtained from ProSci (Cat. 6217); Phospho-c-Jun (Ser73) antibody (Cat. 9164), Phospho-c-Jun (Ser63) II antibody (Cat. 9261), c-Jun (60A8) Rabbit mAb antibody (Cat. 9165), and Vinculin (E1E9V) XP Rabbit mAb antibody (Cat. 13901) were all obtained from Cell Signaling Technology. Anti-coronavirus nucleoprotein OC43 antibody, clone 542-7D (Cat. MAB9012) was from Millipore Sigma; SARS-CoV-2/2019-nCoV Spike/S2 Antibody (Cat. 40590-T62), and HCoV-229E Nucleocapsid Antibody (Cat. 40640-T62) were from Sino Biological; Goat anti-Rabbit IgG (H+L) Highly Cross-Adsorbed Secondary Antibody, Alexa Fluor^TM^ 568 (Cat. A-11036); Goat anti-Mouse IgG (H+L) Cross-Adsorbed Secondary Antibody, Alexa Fluor^TM^ 488 (Cat. A-11001), and Anti-rabbit second antibody (Cat. T2767) were from Thermo Fisher Scientific.

#### mRNA Silencers

pLKO.1-puro shRNA control plasmid (Cat. SHC002), pLKO.1-TRCN0000051135 targeting GLS (Clone ID: NM_014905.2-1441s1c1), and pLKO.1-TRCN0000298987 targeting GLS (Clone ID: 014905.3-1475s21c1) were from Millipore Sigma; and Silencer Select Negative Control No.1 siRNA (Cat. 4390843), Silencer Select GLS siRNA (ID. s5838), and Silencer Select GLS2 siRNA (ID. 5840) were from Thermo Fisher Scientific.

#### Inhibitors

SU1 and BPTES were synthesized by Dr. Scott Ulrich (Ithaca College); UP4 (UPGL00004) (Cat. SML2472), originally developed by Dr. Lee McDermott in collaboration with the Cerione laboratory was obtained from Millipore Sigma; 968 was from ChemBridge Corporation; the c-Jun N-terminal kinase inhibitor SP600125 (Cat. S5567) and the c-Myc inhibitor 10058-F4 (Cat. F3680) were from Millipore Sigma.

#### RNAs isolation kits

Direct-zol RNA MicroPrep was from ZYMO Research (cat. R2062), and the NucleoSpin RNA Virus was from Takara (cat. 740956)

#### qPCR primers and kit

SARS-CoV-2-spike-F (TGGCCGCAAATTGCACAATT)

SARS-CoV-2-spike-R (TGTAGGTCAACCACGTTCCC)

SARS-COV-2 probe (FAM/CGCATTGGCATGGAAGTCAC/BHQ)

HCoV-OC43-F (CCCAAGCAAACTGCTACCTCTCAG)

HCoV-OC43-R (CCCAAGCAAACTGCTACCTCTCAG)

HCoV-229E-F (TCTGCCAAGAGTCTTGCTCG)

HCoV-229E-R (TCTGCCAAGAGTCTTGCTCG)

GLS-F (TGTCACGATCTTGTTTCTCTGTG)

GLS-R (TCATAGTCCAATGGTCCAAAG)

GLS2-F (GCCTGGGTGATTTGCTCTTTT)

GLS2-R (CCTTTAGTGCAGTGGTGAACTT)

actin*-*F (CATCGAGCACGGCATCGTCA)

actin-R (TAGCACAGCCTGGATAGCAAC)

iTaq™ Universal SYBR^®^ Green One-Step Kit was from Bio-Red (cat. 1725151)

## Methods

### SARS-CoV-2 propagation and infection

Studies of SARS-CoV-2 were performed in a BSL3 lab. VeroE6 cells were grown in T75 flasks until 80-90% confluent. SARS-CoV-2 virus dilution was prepared in 2 ml of DMEM 2% FBS (MOI 0.1) per flask. After washing with PBS twice, virus dilution was added to the cell flasks for 1 hour at 37°C in 5%CO_2_ with continuous shaking, followed by adding 10 ml DMEM 2%FBS medium and incubating for 4-6 days or until the cytopathic effect (CPE) progressed through 80% of the cells. The virus medium was collected and centrifuged at 1200 rpm for 10 minutes. The supernatant containing the virus was aliquoted and stored at -80°C in liquid Nitrogen. Plaque assays and qPCR were performed to determine the virus titer.

VeroE6 cells were grown in T75 flasks until 70% confluent, pre-treated with inhibitors for 24 hours, and then infected with SARS-CoV-2 (MOI 0.01) for 1 hour at 37°C 5%CO_2_ with shaking. The plate media was changed to the growth medium with or without inhibitors and the incubation continued for 23 hours at 37°C 5%CO_2._ The viral supernatant medium was inactivated by TRIzol and used for RNA isolation and qPCR. The cells were collected and cell lysates were prepared by adding RIPA buffer and then heat inactivated for Western blot assays.

### HCoV-OC43 propagation and infection

The HCoV-OC43 virus propagation followed the ATCC protocol. HCT8 cells were grown in 15 cm plates until 80-90% confluent. Virus dilution was prepared in 7 ml of RPMI medium containing 2% HS (MOI 0.05). The monolayer cell plates were washed twice with PBS. Virus dilution was adsorbed by cells for 2 hours at 33°C 5%CO_2_ in an incubator while shaking continuously. Ten ml of RPMI containing 2% HS medium were added and continued incubating for 4-6 days at 33°C 5%CO_2_ with shaking. The virus medium was collected when the CPE progressed through 80% of the monolayer and then was centrifuged at 1200 rpm for 10 minutes and aliquots of the supernatant containing the virus were stored at -80°C. The virus titer was determined by plaque assay and qPCR analysis.

HCT8 cells or HBEC cells were grown in 10 cm plates until 70-80% confluent at 37°C 5%CO_2_ and pretreated with inhibitors for 3 hours. HCoVOC43 virus (MOI 0.01, RPMI 2% HS) was used to infect cells for 1 hour at 33°C 5%CO_2_ with shaking. The different conditions of the growth medium of HCT8 cells or HBECs were changed as designed and the cell plates were continued to be incubated at 37°C 5%CO_2_ for 23 hours. The cells and media were collected for Western blot and qPCR analysis, respectively.

### HCoV-229E virus propagation and infection

Following the ATCC protocol for virus propagation MRC5 (EMEM 10% FBS) cells were grown in the 15 cm plates until 80-90% confluent. The virus dilution was prepared in 7 ml of EMEM with 2% FBS medium, MOI 0.05. The cells were adsorbed with a virus dilution at 35°C 5%CO_2_ with shaking for 2 hours. Ten ml of EMEM with 2% FBS medium were added to the plates and incubated for 4-6 days until CPE progressed to 80%. The virus media were collected and centrifuged at 1200 rpm for 10 minutes. The supernatants containing viruses were aliquoted and stored at -80°C. The virus titer was determined by qPCR and plaque assays.

MRC5 cells were grown for 70-80% confluent at 37°C 5%CO2 and were pretreated with the inhibitors for 3 hours. They were then infected with the virus (MOI 0.01, EMEM 2% FBS) dilution for 1 hour at 35°C 5%CO_2_ with shaking. The cell media were changed to the appropriate growth media conditions and the cells were incubated for 23 hours at 37°C 5%CO_2_. The cells and media were collected for further analysis.

### Western blot analysis

Western blot analyses were performed as described^13^. Briefly, the cells were collected following different treatment conditions and lysed with lysis buffer. Protein concentrations were determined by Bradford assay (Bio-Rad), and lysate proteins were denatured by boiling for 5 min. Lysate proteins were resolved on Tris-glycine protein gels (Life Technologies) and then transferred to PVDF membranes (PerkinElmer). Membranes were blocked with milk (5%) or BSA (10%) in TBST for at least 1 hour and incubated with primary antibody dilutions in TBST overnight at 4°C. Horseradish-peroxidase-conjugated secondary antibodies were applied to detect primary antibodies, followed by imaging with Western Lighting Plus-ECL (PerkinElmer).

### Immunofluorescence staining

HBECs were grown in the 4-well slide chamber, and diluted HCoV-OC43 viruses (MOI 0.01) were adsorbed for 1 hour at 33°C 5%CO_2_. The culture medium was changed to growth medium and the cells were maintained at 37°C 5%CO_2_ for 22 hours. One drop of NucBlue Live Cell staining solution was added to each cell well for 1 hour. The cells were fixed with 3.7% formaldehyde for 20 minutes and washed with PBS (3X). GLS antibody (1:500) was added to the cell chamber at room temperature for 2 hours, the cells were washed with PBS (3X), and then HCoV-OC43 antibody (1:500) was added to the chamber for 2 hours at room temperature. The mixture of Goat anti-rabbit secondary antibody, Alexa Fluor 568 (1:200) for GLS primary antibody, Goat anti-mouse secondary antibody, and Alexa Fluor 488 (1:200) for HCoV-OC43 primary antibody, were applied to the chamber and incubated at room temperature for 1 hour with rocking. The cells were washed with PBS (3X). The images were taken with a KEYENCE BZ-X810 microscope.

### RNA isolation from cells and media, and quantitative PCR (qPCR)

Total RNA was isolated from cells or media using a Direct-zol RNA MicroPrep kit or a NucleoSpin RNA Virus kit, respectively. qPCR analysis was carried out with specific primers, and iTaq™ Universal SYBR^®^ Green Two-step for total RNA from cells and One-Step for RNA from the media. Reactions were performed using the real-time PCR system (ViiA7 applied biosystems).

### Metabolite Extraction

HBECs were grown in 6-well plates at 80% confluence (triplets for each condition), pretreated with SU1 (2.5μM) or UP4 (0.5μM) for 3 hours with non-treated wells as controls, and then infected with HCoV-OC43 (MOI 0.01) for 1 hour at 33°C 5%CO_2_ with shaking, followed by the media being changed to growth media with or without inhibitors, and the incubations then continued at 37°C 5%CO_2_ for 23 hours. The cell plates were placed on dry ice, and 1 ml of ice-cold extraction solution was added to each well of the cells with the cells remaining on dry ice for 10 minutes. The cells were scraped in the extraction solution and transferred to Eppendorf tubes. The tubes were incubated on dry ice for 1 hour and then centrifuged at 13000 rpm for 10 minutes. Supernatants (700μl) from each sample were transferred to new tubes, and then samples were analyzed at the Cold Spring Harbor Metabolomic Facility.

### Metabolomics Analysis

Metabolite levels were determined for targeted metabolomics analysis of HBECs that were either uninfected, HCoV-OC43-infected, or HCoV-OC43-infected and treated with UP4 or SU1. The heatmaps of different metabolic pathways were produced using the Cluster Analysis module under the same platform using the metabolite levels in individual pathways as inputs with the same parameters.

### Genetic Knockdowns using shRNA and siRNA

Knockdowns of GLS expression in HBECs cells were achieved using short hairpin RNA (shRNA). Lentivirus particles for each shRNA construct were generated using exponentially growing 293T cells (ATCC) as described previously^11,21^. Silencer Select pre-designed siRNAs targeting GLS and the control silencer were transfected into HBECs, using 60 mm dishes, with 0.3 ml Opti-MEM(GIBCO) containing 100nM of the appropriate siRNA to give the final concentration of 10nM, along with 0.3 ml Opti-MEM containing 12 µl of Lipofectamine 2000 (Invitrogen), were incubated separately at room temperature for 5 minutes. The two solutions were then combined and incubated for an additional 20 minutes, mixed with 2.4 ml culture medium, and added to cells. After 5 hours of incubation at 37°C, the transfection mixture was replaced with fresh culture medium. For all the knockdowns, two independent shRNAs or siRNAs were used, along with negative control shRNA or siRNA.

### Virus plaque assay

A total of 2X10^5^ host cells were seeded in 6-well plates with culture medium and incubated for 24 hours at 37°C. The wells were washed with PBS, and then the cells were infected with mock supernatant or different dilutions of virus (100µl), in triplicate per condition. The plates were incubated at 37°C for 4 hours and mixed gently every 15 minutes. Sterile 0.4% Oxoid ager in culture medium was prepared and 2 ml was applied to each well to replace the medium. The cell plates were placed in a tissue culture hood for 15 minutes as the agar overlay turned solid, then incubated at 37°C 5%CO_2_ for 5-7 days. PFA (4%) was used to fix the cells for 30 minutes at room temperature, and then the agar was removed. Following immunostaining with virus antibody or Crystal violet staining, plaques were counted.

## Statistical analyses

All differences were analyzed with one-way ANOVA with Bonferroni correction and each experiment was repeated independently at least three times.

## Acknowledgments

We thank Dr. Scott Ulrich (Ithaca College) for designing and synthesizing SU1 compound, and Dr. Lee McDermott (formerly of the University of Pittsburgh) for developing UP4 compound with us. We also thank the Metabolomic Facility at Cold Spring Harbor for the metabolomic assay.

## Author contribution

K.S.G, K.F.W and R.A.C designed experiments. K.S.G, A.C, M.C, N.Y, R.L, Y.Q, K.S.R, J.S and W.P.K performed the experiments and data analysis. K.S.G, Y.Q and W.P.K prepared the figures. K.S.G, M.J.L, G.R.W and R.A.C contributed to the writing and editing of the manuscript.

## Supplementary Figures

**Figure S1.**
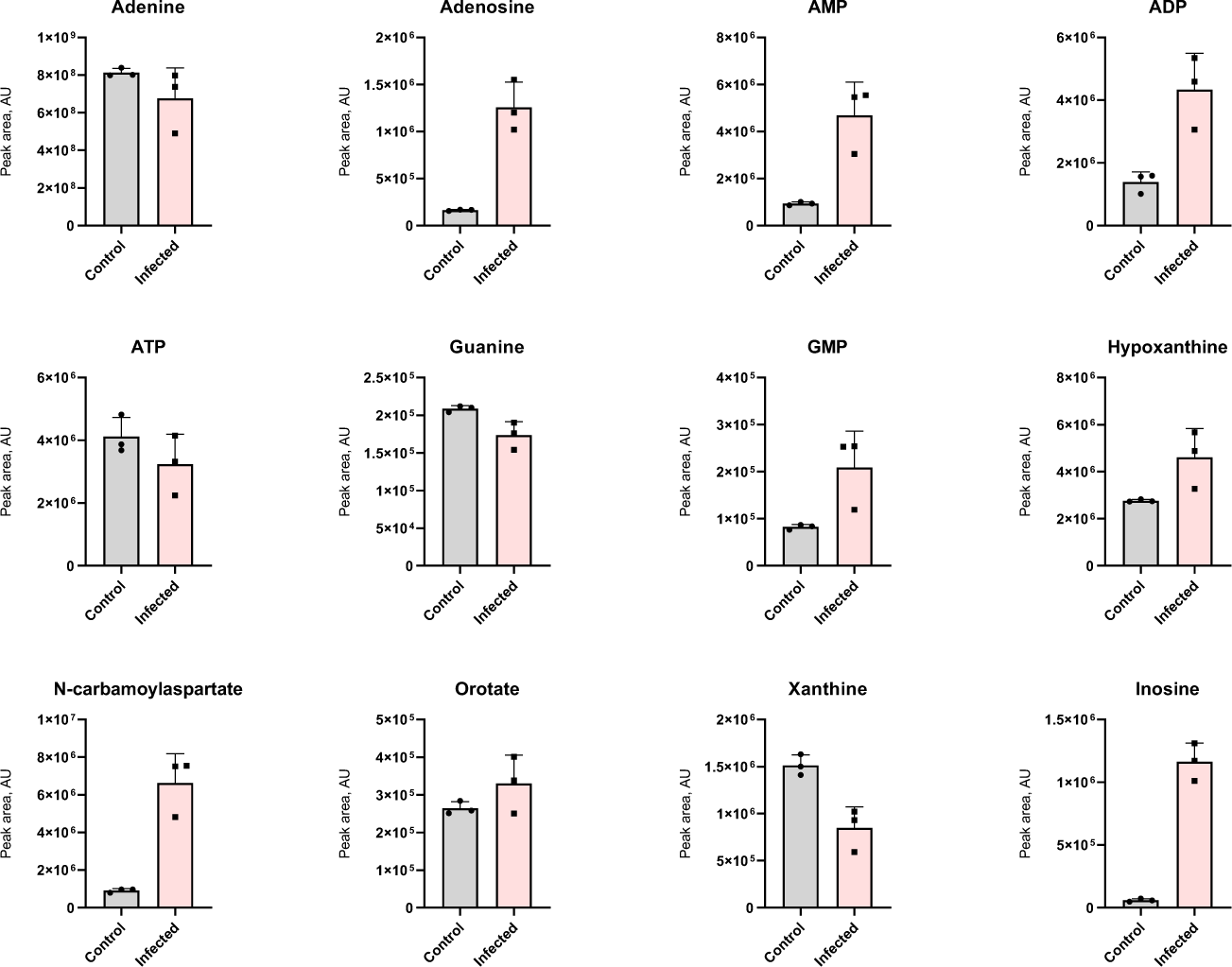
Metabolomic data—Purine axis. Bar graphs from metabolomics showed the metabolite changes in the purine synthetic pathway in HBECs infected with HCoV-OC43 compared to uninfected samples. Data are presented as mean +/-SD (n=3 independent biological replicates).

**Figure S2.**
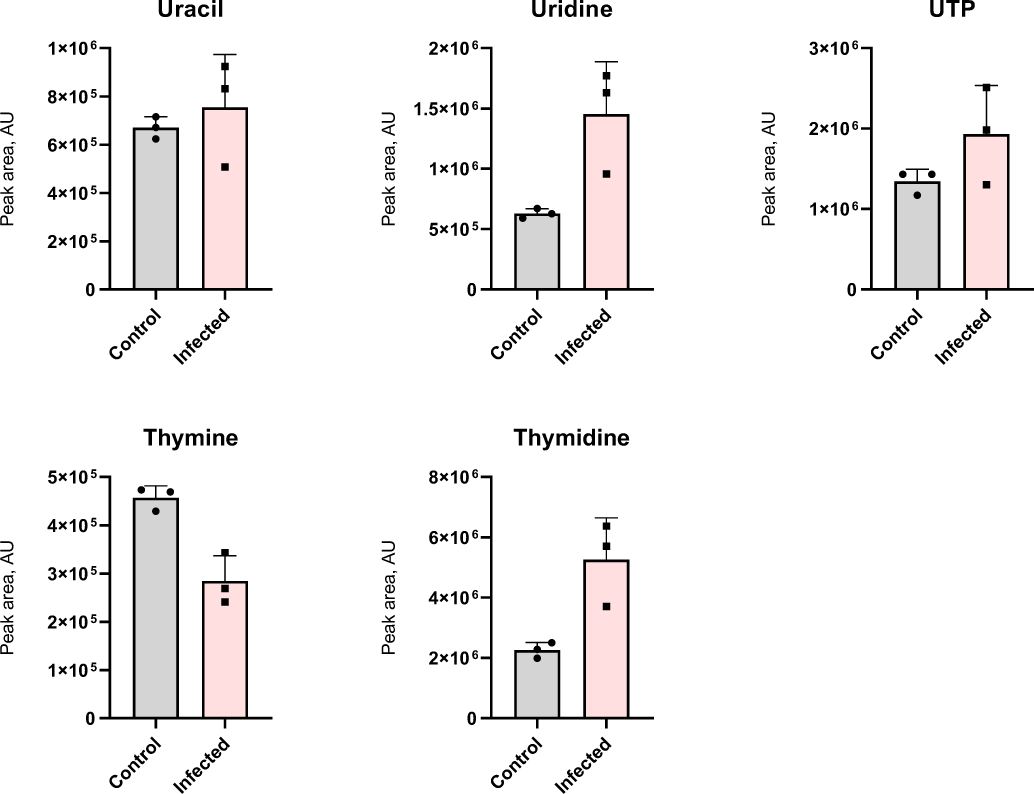
Metabolomic data—Pyrimidine axis. Bar graphs from metabolomics showed the metabolite changes in the pyrimidine synthetic pathway in HBECs infected with HCoV-OC43 compared to uninfected samples. Data are presented as mean +/-SD (n=3 independent biological replicates).

**Figure S3.**
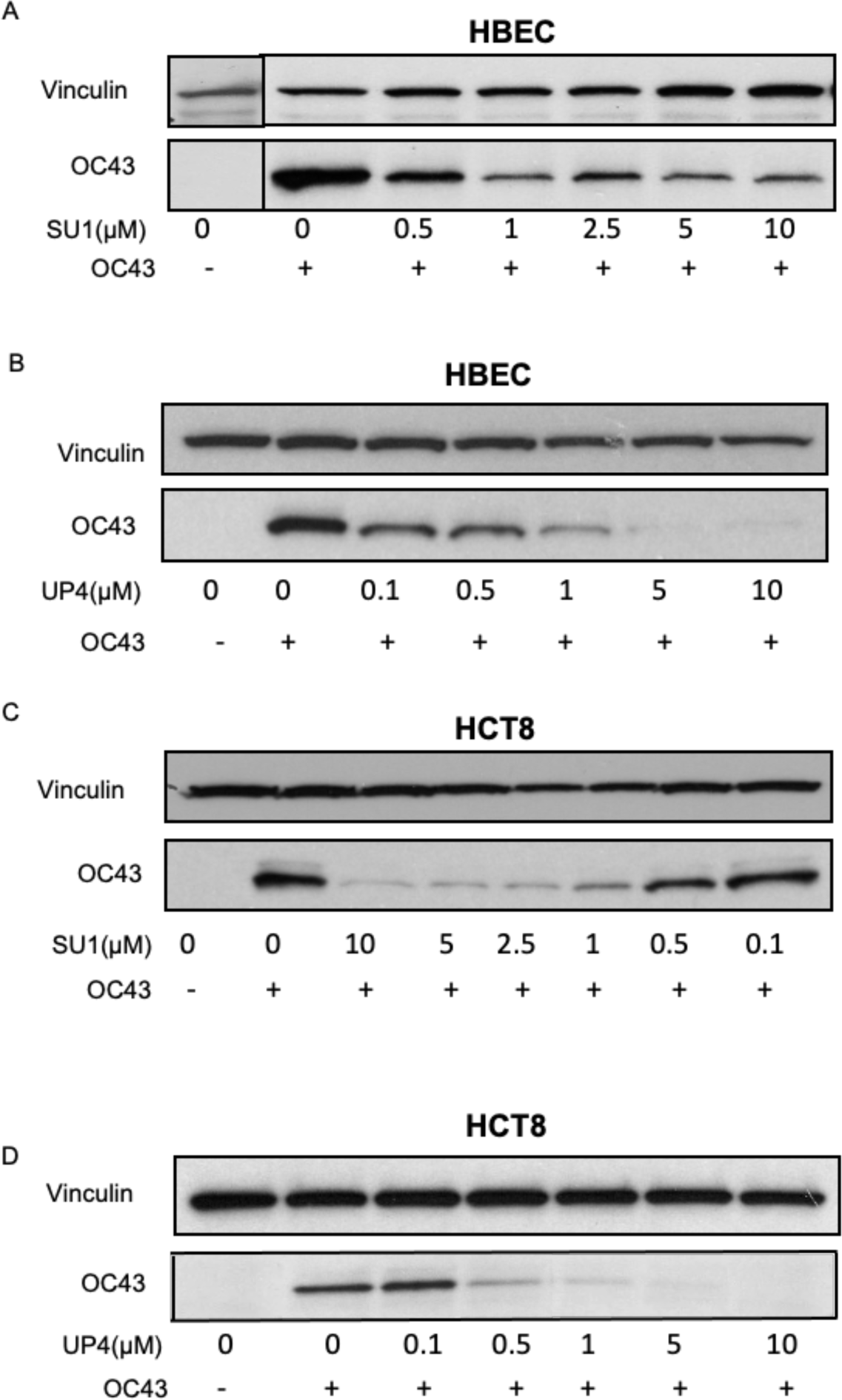
Dose-dependent inhibition of virus-infected HBECs and HCT8 cells by the SU1 and UP4 compounds. (A) HBECs were pretreated with different doses of SU1 (0, 0.5, 1, 2.5, 5, 10mM) for 3 hours, followed by infection with HCoV-OC43 (MOI 0.01, 2%HS RPMI, 33° C 5%CO_2_ for 1 hour). The cells were changed to the growth media with different concentrations of SU1 and were incubated for 23 hours at 37°C 5%CO_2_. Western blot analysis showed the effects of SU1 on HCoV-OC43 replication in HBECs. (B) A similar experiment was performed to examine the effects of the UP4 compound. Western blot analysis showed the effects of varying doses of UP4 (0, 0.1, 0.5, 1, 5, 10mM) upon a 24-hour virus infection of HBECs. (C) HCT8 cells were pretreated with different concentrations of SU1 for 3 hours and infected with HCoV-OC43, as described above. Western blot analysis showed the different virus levels as a function of SU1 concentrations. (D) HCT8 cells were pretreated with varying doses of UP4 compound and then infected with HCoV-OC43. Western blot analysis showed the HCoV-OC43 replication levels as a function of different amounts of UP4.

**Figure S4.**
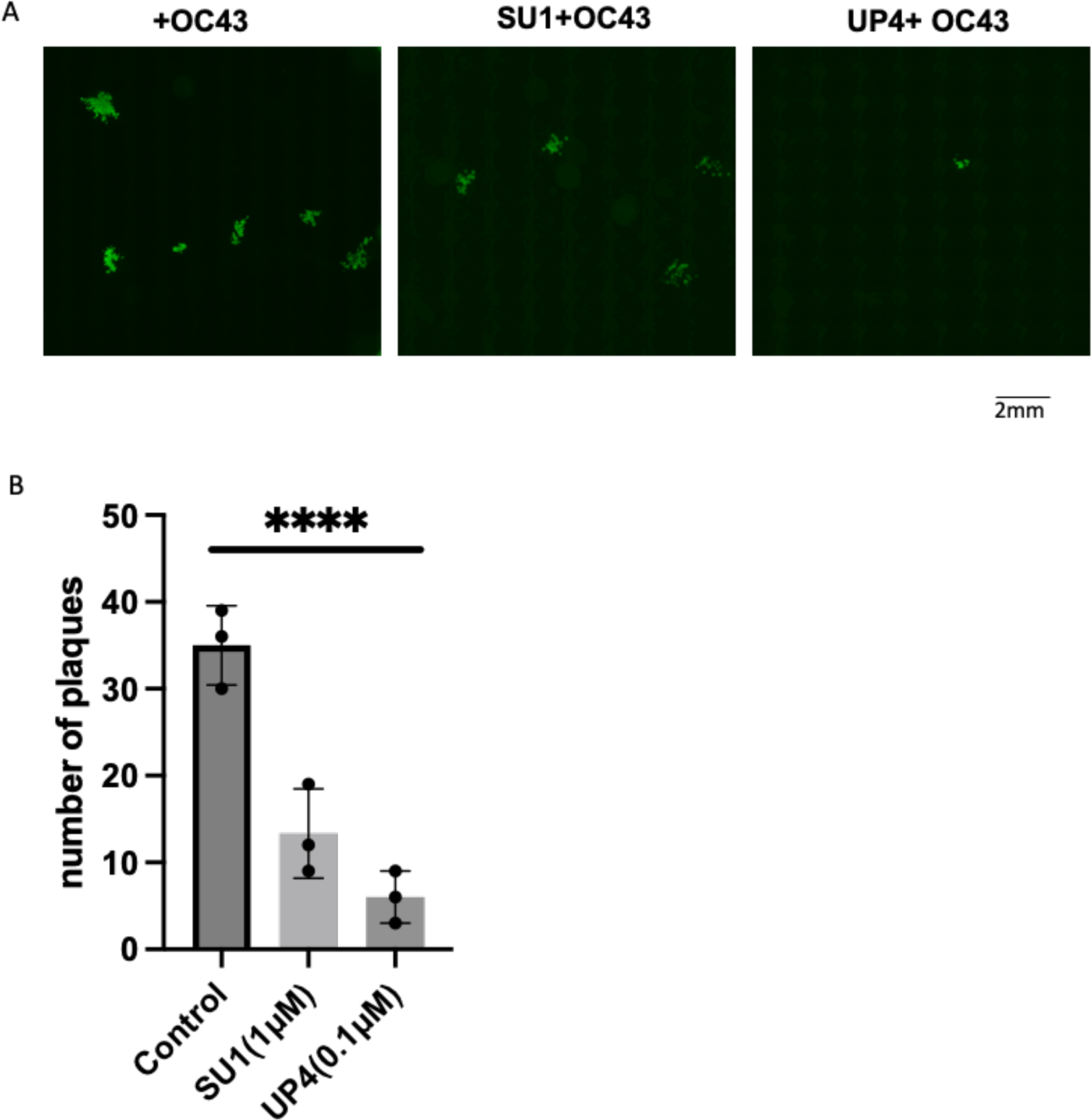
SU1 and UP4 inhibit HCoV-OC43 replication in HBECs by plaque assays. HBECs (2X10^5^) were seeded in 6-well plates, pretreated with DMSO, SU1(2.5µM), and UP4(0.1µM) for 3 hours, infected with HCoV-OC43 (MOI 0.01) at 33°C 5%CO_2_ with shaking for 1 hour, at which point the agar medium was replaced with the virus solution after a PBS wash. The plates were maintained at room temperature hood for 15 minutes and then incubated at 37°C 5%CO_2_ for 4 days. The agar layer was removed, and cells were fixed with 4% PFA and stained with HCoV-OC43 antibody and goat anti-mouse secondary antibody. Triplicate determinations were performed for each condition. (A) imaging of the plaques. (B) The bar graph shows the relative plaque numbers for virus-infected cells treated with SU1 or UP4, DMSO control-treated cells. One-way ANOVA with Bonferroni correction was used to determine significance in B, **** indicates modified *P* < 0.0001.

**Figure S5.**
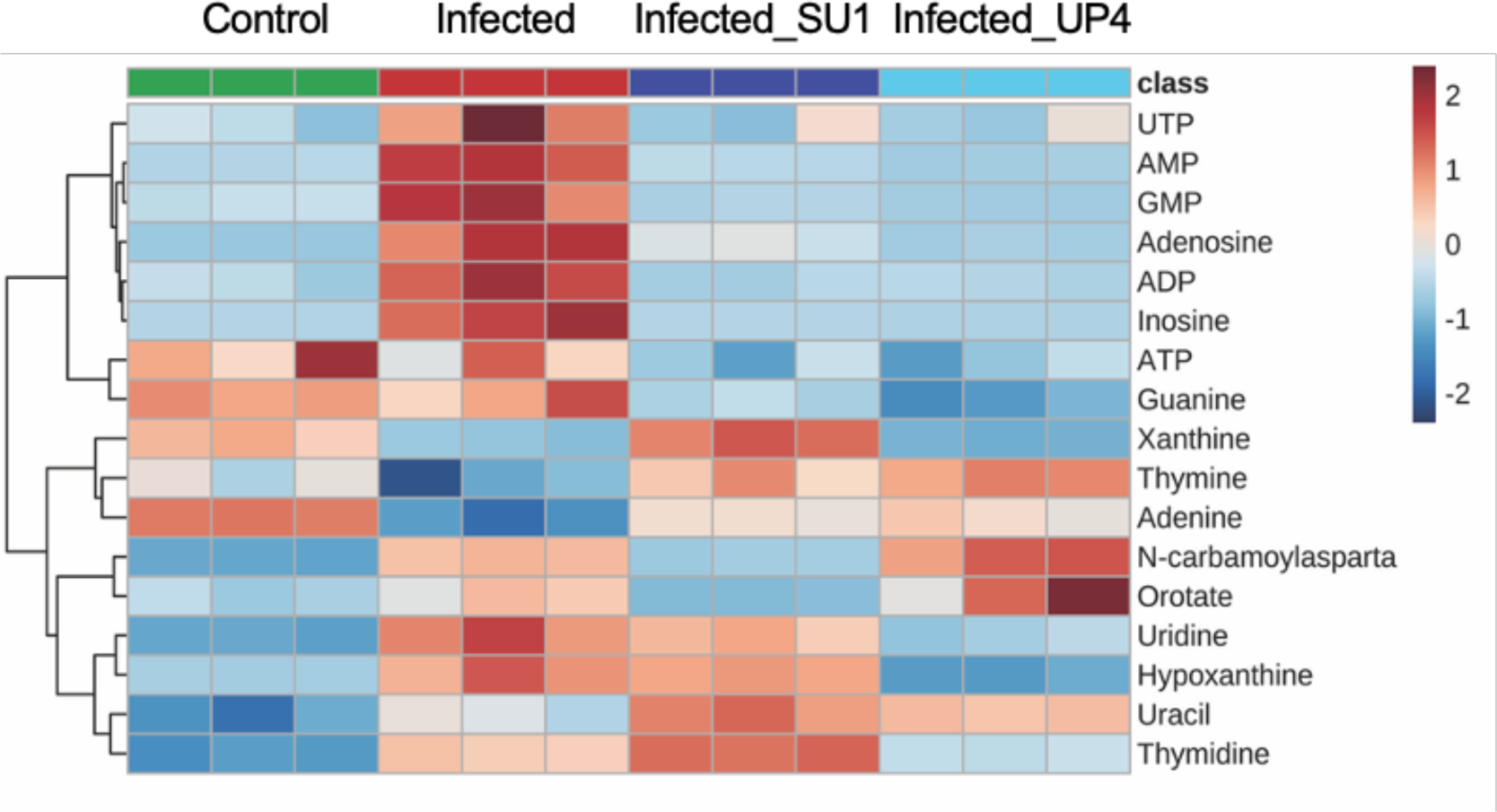
Coronavirus infection affects nucleotide synthesis in host cells. Heatmap for metabolites in uninfected HBECs, HBECs infected with HCoV-OC43, and infected HBECs treated with SU1(2.5µM) and UP4(0.5µM).

## REFERENCES

1. V’kovski P, Kratzel A, Steiner S, Stalder H, Thiel V. Coronavirus biology and replication: implications for SARS-CoV-2. Nat Rev Microbiol. 2021;19(3):155–170. doi:10.1038/s41579-020-00468-6

2. Zhou P, Yang X-L, Wang X-G, et al. A pneumonia outbreak associated with a new coronavirus of probable bat origin. Nature. 2020;579(7798):270–273. doi:10.1038/s41586-020-2012-7

3. Beyerstedt S, Casaro EB, Rangel ÉB. COVID-19: angiotensin-converting enzyme 2 (ACE2) expression and tissue susceptibility to SARS-CoV-2 infection. Eur J Clin Microbiol Infect Dis. 2021;40(5):905–919. doi:10.1007/s10096-020-04138-6

4. Shapira T, Monreal IA, Dion SP, et al. A TMPRSS2 inhibitor acts as a pan-SARS-CoV-2 prophylactic and therapeutic. Nature. 2022;605(7909):340–348. doi:10.1038/s41586-022-04661-w

5. Liu DX, Liang JQ, Fung TS. Human Coronavirus-229E, -OC43, -NL63, and - HKU1 (Coronaviridae). In: Encyclopedia of Virology. Elsevier; 2021:428–440. doi:10.1016/B978-0-12-809633-8.21501-X

6. Sriwilaijaroen N, Suzuki Y. Host receptors of influenza viruses and coronaviruses—molecular mechanisms of recognition. Vaccines. 2020;8(4):1–47. doi:10.3390/vaccines8040587

7. Tang AT, Buchholz DW, Szigety KM, et al. Cell-autonomous requirement for ACE2 across organs in lethal mouse SARS-CoV-2 infection. PLOS Biol. 2023;21(2):e3001989. doi:10.1371/journal.pbio.3001989

8. Mayer KA, Stöckl J, Zlabinger GJ, Gualdoni GA. Hijacking the supplies: Metabolism as a novel facet of virus-host interaction. Front Immunol. 2019;10(JULY):1–12. doi:10.3389/fimmu.2019.01533

9. Fontaine KA, Camarda R, Lagunoff M. Vaccinia Virus Requires Glutamine but Not Glucose for Efficient Replication. J Virol. 2014;88(8):4366–4374. doi:10.1128/jvi.03134-13

10. Pavlova NN, Thompson CB. The Emerging Hallmarks of Cancer Metabolism. Cell Metab. 2016;23(1):27–47. doi:10.1016/j.cmet.2015.12.006

11. De Berardinis RJ, Chandel NS. Fundamentals of cancer metabolism. Sci Adv. 2016;2(5). doi:10.1126/sciadv.1600200

12. Finley LWS. What is cancer metabolism ? 2023;(March):2023.

13. Greene KS, Lukey MJ, Wang X, et al. SIRT5 stabilizes mitochondrial glutaminase and supports breast cancer tumorigenesis. Proc Natl Acad Sci U S A. 2019;116(52):26625–26632. doi:10.1073/pnas.1911954116

14. Vander Heiden MG, DeBerardinis RJ. Understanding the Intersections between Metabolism and Cancer Biology. Cell. 2017;168(4):657–669. doi:10.1016/j.cell.2016.12.039

15. Martı M, Cardona C, Rosa V De, et al. Neurochemistry International A novel glutaminase isoform in mammalian tissues. 2009;55:76–84. doi:10.1016/j.neuint.2009.02.021

16. Altman BJ, Stine ZE, Dang C V. From Krebs to Clinic : Glutamine Metabolism to Cancer Therapy. Nat Rev Cancer. 2017;16(10):619–634. doi:10.1038/nrc.2016.71.From

17. Ferrer CM, Lynch TP, Sodi VL, et al. O-GlcNAcylation Regulates Cancer Metabolism and Survival Stress Signaling via Regulation of the HIF-1 Pathway. Mol Cell. 2014;54(5):820–831. doi:10.1016/j.molcel.2014.04.026

18. Katt WP, Lukey MJ, Cerione RA. A tale of two glutaminases: Homologous enzymes with distinct roles in tumorigenesis. Future Med Chem. 2017;9(2):223–243. doi:10.4155/fmc-2016-0190

19. Campos-Sandoval JA, Martín-Rufián M, Cardona C, Lobo C, Peñalver A, Márquez J. Glutaminases in brain: Multiple isoforms for many purposes. Neurochem Int. 2015;88:1–5. doi:10.1016/j.neuint.2015.03.006

20. Wang J Bin, Erickson JW, Fuji R, et al. Targeting mitochondrial glutaminase activity inhibits oncogenic transformation. Cancer Cell. 2010;18(3):207–219. doi:10.1016/j.ccr.2010.08.009

21. Lukey MJ, Cluntun AA, Katt WP, et al. Liver-Type Glutaminase GLS2 Is a Druggable Metabolic Node in Luminal-Subtype Breast Cancer. Cell Rep. 2019;29(1):76–88.e7. doi:10.1016/j.celrep.2019.08.076

22. Gao P, Tchernyshyov I, Chang TC, et al. C-Myc suppression of miR-23a/b enhances mitochondrial glutaminase expression and glutamine metabolism. Nature. 2009;458(7239):762–765. doi:10.1038/nature07823

23. Lukey MJ, Greene KS, Erickson JW, Wilson KF, Cerione RA. The oncogenic transcription factor c-Jun regulates glutaminase expression and sensitizes cells to glutaminase-targeted therapy. Nat Commun. 2016;7:1–14. doi:10.1038/ncomms11321

24. Dias MM, Adamoski D, dos Reis LM, et al. GLS2 is protumorigenic in breast cancers. Oncogene. 2020;39(3):690–702. doi:10.1038/s41388-019-1007-z

25. Milano SK, Huang Q, Nguyen TTT, et al. New insights into the molecular mechanisms of glutaminase C inhibitors in cancer cells using serial room temperature crystallography. J Biol Chem. 2022;298(2):101535. doi:10.1016/j.jbc.2021.101535

26. Wicker CA, Hunt BG, Krishnan S, et al. Glutaminase inhibition with telaglenastat (CB-839) improves treatment response in combination with ionizing radiation in head and neck squamous cell carcinoma models. Cancer Lett. 2021;502:180–188. 10.1016/j.canlet.2020.12.038

27. Gross MI, Demo SD, Dennison JB, et al. Antitumor Activity of the Glutaminase Inhibitor CB-839 in Triple-Negative Breast Cancer. Mol Cancer Ther. 2014;13(4):890–901. doi:10.1158/1535-7163.MCT-13-0870

28. Bharadwaj S, Singh M, Kirtipal N, Kang SG. SARS-CoV-2 and Glutamine: SARS-CoV-2 Triggered Pathogenesis via Metabolic Reprograming of Glutamine in Host Cells. Front Mol Biosci. 2021;7. doi:10.3389/fmolb.2020.627842

29. Thaker SK, Ch’ng J, Christofk HR. Viral hijacking of cellular metabolism. BMC Biol. 2019;17(1):59. doi:10.1186/s12915-019-0678-9

30. Thai M, Thaker SK, Feng J, et al. MYC-induced reprogramming of glutamine catabolism supports optimal virus replication. Nat Commun. 2015;6(1):8873. doi:10.1038/ncomms9873

31. Schultz DC, Johnson RM, Ayyanathan K, et al. Pyrimidine inhibitors synergize with nucleoside analogues to block SARS-CoV-2. Nature. 2022;604(7904):134–140. doi:10.1038/s41586-022-04482-x

32. Chen J, Ye C, Wan C, et al. The Roles of c-Jun N-Terminal Kinase (JNK) in Infectious Diseases. Int J Mol Sci. 2021;22(17):9640. doi:10.3390/ijms22179640

33. Mizutani T, Fukushi S, Saijo M, Kurane I, Morikawa S. JNK and PI3k/Akt signaling pathways are required for establishing persistent SARS-CoV infection in Vero E6 cells. Biochim Biophys Acta - Mol Basis Dis. 2005;1741(1-2):4–10. doi:10.1016/j.bbadis.2005.04.004

34. Varshney B, Lal SK. SARS-CoV Accessory Protein 3b Induces AP-1 Transcriptional Activity through Activation of JNK and ERK Pathways. Biochemistry. 2011;50(24):5419–5425. doi:10.1021/bi200303r

35. Leite FGG, Torres AA, De Oliveira LC, et al. c-Jun integrates signals from both MEK/ERK and MKK/JNK pathways upon vaccinia virus infection. Arch Virol. 2017;162(10):2971–2981. doi:10.1007/s00705-017-3446-6

36. McDermott LA, Iyer P, Vernetti L, et al. Design and evaluation of novel glutaminase inhibitors. Bioorg Med Chem. 2016;24(8):1819–1839. doi:10.1016/j.bmc.2016.03.009

37. Katt WP, Ramachandran S, Erickson JW, Cerione RA. Dibenzophenanthridines as inhibitors of glutaminase C and cancer cell proliferation. Mol Cancer Ther. 2012;11(6):1269–1278. doi:10.1158/1535-7163.MCT-11-0942

38. Stalnecker C a, Ulrich SM, Li Y, et al. Mechanism by which a recently discovered allosteric inhibitor blocks glutamine metabolism in transformed cells. Proc Natl Acad Sci U S A. 2014;112(2):394–399. doi:10.1073/pnas.1414056112

39. Robinson MM, Mcbryant SJ, Tsukamoto T, et al. Novel mechanism of inhibition of rat kidney-type glutaminase by bis-2-(5-phenylacetamido-1,2,4-thiadiazol-2-yl)ethyl sulfide (BPTES). Biochem J. 2007;406(3):407–414. doi:10.1042/BJ20070039

40. Huang Q, Stalnecker C, Zhang C, et al. Characterization of the interactions of potent allosteric inhibitors with glutaminase C, a key enzyme in cancer cell glutamine metabolism. J Biol Chem. 2018;293(10):3535–3545. doi:10.1074/jbc.M117.810101

41. Best SA, Gubser PM, Sethumadhavan S, et al. Glutaminase inhibition impairs CD8 T cell activation in STK11-/Lkb1-deficient lung cancer. Cell Metab. 2022;34(6):874–887.e6. doi:10.1016/j.cmet.2022.04.003

42. Sanchez EL, Pulliam TH, Dimaio TA, Thalhofer AB, Delgado T, Lagunoff M. Glycolysis, Glutaminolysis, and Fatty Acid Synthesis Are Required for Distinct Stages of Kaposi’s Sarcoma-Associated Herpesvirus Lytic Replication. J Virol. 2017;91(10). doi:10.1128/JVI.02237-16

43. DeBerardinis RJ, Lum JJ, Hatzivassiliou G, Thompson CB. The Biology of Cancer: Metabolic Reprogramming Fuels Cell Growth and Proliferation. Cell Metab. 2008;7(1):11–20. doi:10.1016/j.cmet.2007.10.002

44. Jiang B, Zhang J, Zhao G, et al. Filamentous GLS1 promotes ROS-induced apoptosis upon glutamine deprivation via insufficient asparagine synthesis. Mol Cell. 2022;82(10):1821–1835.e6. doi:10.1016/j.molcel.2022.03.016

45. Feng S, Aplin C, Nguyen TT, Milano SK, Cerione RA. Filament Formation drives catalysis by glutaminase enzymes important in cancer progression. bioRxiv. July 2023.

46. Varghese S, Pramanik S, Williams LJ, et al. The glutaminase inhibitor CB-839 (Telaglenastat) enhances the antimelanoma activity of T-cell-mediated immunotherapies. Mol Cancer Ther. 2021;20(3):500–511. doi:10.1158/1535-7163.MCT-20-0430

47. McDermott L, Koes D, Mohammed S, et al. GAC inhibitors with a 4-hydroxypiperidine spacer: Requirements for potency. Bioorg Med Chem Lett. 2019;29(19):126632. doi:10.1016/j.bmcl.2019.126632

48. Harding JJ, Telli M, Munster P, et al. A phase I dose-escalation and expansion study of telaglenastat in patients with advanced or metastatic solid tumors. Clin Cancer Res. 2021;27(18):4994–5003. doi:10.1158/1078-0432.CCR-21-1204

49. Riess JW, Frankel P, Shackelford D, et al. Phase 1 Trial of MLN0128 (Sapanisertib) and CB-839 HCl (Telaglenastat) in Patients With Advanced NSCLC (NCI 10327): Rationale and Study Design. Clin Lung Cancer. 2021;22(1):67–70. doi:10.1016/j.cllc.2020.10.006

50. Seifert M, Bera SC, Van Nies P, et al. Inhibition of sars-cov-2 polymerase by nucleotide analogs from a single-molecule perspective. Elife. 2021;10. doi:10.7554/eLife.70968

51. Mouffouk C, Mouffouk S, Mouffouk S, Hambaba L, Haba H. Flavonols as potential antiviral drugs targeting SARS-CoV-2 proteases (3CLpro and PLpro), spike protein, RNA-dependent RNA polymerase (RdRp) and angiotensin-converting enzyme II receptor (ACE2). Eur J Pharmacol. 2021;891:173759. doi:10.1016/j.ejphar.2020.173759

52. Tian L, Qiang T, Liang C, et al. RNA-dependent RNA polymerase (RdRp) inhibitors: The current landscape and repurposing for the COVID-19 pandemic. Eur J Med Chem. 2021;213:113201. doi:10.1016/j.ejmech.2021.113201

53. Fernandes Q, Inchakalody VP, Merhi M, et al. Emerging COVID-19 variants and their impact on SARS-CoV-2 diagnosis, therapeutics and vaccines. Ann Med. 2022;54(1):524–540. doi:10.1080/07853890.2022.2031274

54. Peng F, Yuan H, Wu S, Zhou Y. Recent Advances on Drugs and Vaccines for COVID-19. *Inq J Heal Care Organ Provision*, Financ. 2021;58:004695802110556. doi:10.1177/00469580211055630

55. Bai Z, Cao Y, Liu W, Li J. The SARS-CoV-2 Nucleocapsid Protein and Its Role in Viral Structure, Biological Functions, and a Potential Target for Drug or Vaccine Mitigation. Viruses. 2021;13(6):1115. doi:10.3390/v13061115

56. Pavlova NN, Thompson CB. The Emerging Hallmarks of Cancer Metabolism. Cell Metab. 2016;23(1):27–47. doi:10.1016/j.cmet.2015.12.006

57. Bharadwaj S, Singh M, Kirtipal N, Kang SG. SARS-CoV-2 and Glutamine: SARS-CoV-2 Triggered Pathogenesis via Metabolic Reprograming of Glutamine in Host Cells. Front Mol Biosci. 2021;7(January):1–14. doi:10.3389/fmolb.2020.627842

58. Ferreira APS, Cassago A, Gonçalves K de A, et al. Active Glutaminase C Self-assembles into a Supratetrameric Oligomer That Can Be Disrupted by an Allosteric Inhibitor. J Biol Chem. 2013;288(39):28009–28020. doi:10.1074/jbc.M113.501346

59. Møller M, Nielsen SS, Ramachandran S, et al. Small Angle X-Ray Scattering Studies of Mitochondrial Glutaminase C Reveal Extended Flexible Regions, and Link Oligomeric State with Enzyme Activity. PLoS One. 2013;8(9):e74783. doi:10.1371/journal.pone.0074783

